# Evaluating the Probability of CRISPR-based Gene Drive Contaminating Another Species

**DOI:** 10.1101/776609

**Authors:** Virginie Courtier-Orgogozo, Antoine Danchin, Pierre-Henri Gouyon, Christophe Boëte

## Abstract

The probability D that a given CRISPR-based gene drive element contaminates another, non-target species can be estimated by the following Drive Risk Assessment Quantitative Estimate (DRAQUE) Equation:

D = (*hyb+transf).express.cut.flank.immune.nonextinct* **with**
*hyb* = probability of hybridization between the target species and a non-target species
*transf* = probability of horizontal transfer of a piece of DNA containing the gene drive cassette from the target species to a non-target species (with no hybridization)
*express* = probability that the *Cas9* and guide RNA genes are expressed
*cut* = probability that the CRISPR-guide RNA recognizes and cuts at a DNA site in the new host
*flank* = probability that the gene drive cassette inserts at the cut site
*immune* = probability that the immune system does not reject *Cas9*-expressing cells
*nonextinct* = probability of invasion of the drive within the population

We discuss and estimate each of the seven parameters of the equation, with particular emphasis on possible transfers within insects, and between rodents and humans. We conclude from current data that the probability of a gene drive cassette to contaminate another species is not insignificant. We propose strategies to reduce this risk and call for more work on estimating all the parameters of the formula.

## Introduction

Selfish genetic elements are biological entities that favor their own transmission across generations. Examples include transposons that insert copies of themselves at other places in the genome, homing endonuclease genes that copy themselves at targeted genomic sites, segregation distorters that destroy competing chromosomes during meiosis and maternally heritable microorganisms such as *Wolbachia* that favor progeny of infected females (Agren and Clark 2018). In recent years, researchers have started to develop CRISPR-based gene drives, named here gene drive for short, with the intention to spread synthetic genetic elements into wild populations. Potential applications of gene drives are numerous and include the elimination of mosquitoes to fight malaria, Zika, and other mosquito-borne diseases, or alternatively the modification of mosquitoes from vector to non-vector so that they no longer transmit human pathogens (Esvelt *et al.* 2014). Applications are not restricted to public health issues and also include agriculture, with for instance the elimination of invasive and pest species such as *Drosophila suzukii* or the suppression of herbicide resistance in weeds (Scott *et al.* 2018). Potential uses of gene drive are also reaching the field of conservation biology, with the targeting of rats (*Rattus rattus* and *Rattus norvegicus)* in New Zealand (Leitschuh *et al.* 2018; Rode *et al.* 2019). So far, CRISPR-based gene drives have only been tested in laboratories or in large indoor cages. They have been shown to efficiently boost their own transmission in yeasts (DiCarlo *et al.* 2015), Drosophila flies (Gantz and Bier 2015; Champer *et al.* 2017; KaramiNejadRanjbar *et al.* 2018), mosquitoes (Gantz *et al.* 2015; Hammond *et al.* 2016; Kyrou *et al.* 2018), the pathogenic fungus *Candida albicans* (Shapiro *et al.* 2018) and mice (Grunwald *et al.* 2019).

A CRISPR gene drive cassette is a piece of DNA that comprises several elements: (i) a gene encoding a guide RNA (gRNA) that can recognize a specific target DNA sequence, (ii) a *Cas9* gene encoding a Cas9 endonuclease that can cut DNA at the site specified by the gRNA, (iii) sequences at the extremities that are homologous to sequences flanking the target site, so that the gene drive cassette can copy itself at the cleavage site via homology-directed repair and (iv) optional sequences, for example conferring a trait of interest such as malaria resistance (Esvelt *et al.* 2014). By converting heterozygotes for the gene drive allele into homozygotes, the gene drive cassette alters Mendelian transmission and can thus spread into wild populations. The release in the wild of a few individuals carrying gene drive constructs is thus expected to be sufficient to transform an entire population after a dozen generations (Deredec *et al.* 2008). Gene drives can be designed to introduce a phenotype of interest in a targeted population either through the introduction of a new gene, or by the inactivation of an endogenous gene via the insertion of the gene drive cassette into it (Esvelt *et al.* 2014). With introduced genetic changes that decrease viability or fertility, a gene drive can be used to eradicate a targeted population or to reduce its size, while with other types of genetic changes it is possible to alter the characteristics of a population.

The possibilities offered to humanity in terms of benefits by the new molecular techniques of genome edition CRISPR-Cas are innumerable but also associated with risks which should be carefully monitored (Zhang 2019). One obvious risk associated with gene drive is that the sequence may escape from the target species and spread into other species. Such spillover could have devastating effects, such as the extinction of a species, or the modification of a large number of individuals, with potentially important ecological consequences. Compared to natural bacterial CRISPR systems, gene drive cassettes are more compact and contain eukaryotic cis-regulatory elements, so that they are one step closer to potential contamination of non-target eukaryote species. The risk of gene drives contaminating another species has been mentioned by several authors (Benedict *et al.* 2008; Esvelt *et al.* 2014; Webber *et al.* 2015; National Academies of Sciences and Medicine 2016; Rode *et al.* 2019) but to our knowledge it has not been examined in detail. Risk assessment studies classically present the value of a risk as a product of two terms: Risk = Probability of occurrence x Damage in case of occurrence. The aim of this paper is to derive a formula to evaluate the probability of occurrence of a drive sequence escaping from the target species and endangering another species. The damage resulting from such an unwanted event will obviously depend on the concerned species. If it is simply another mosquito, the damage will be limited (or this could even be seen as a serendipitous positive externality). If it were a keystone species or humans for instance, damages could be very important and hard, not to say impossible to mitigate. This paper does not address these questions but concentrates on the probability that a drive escapes from the target species and contaminates another species. Such an event results from a succession of events and the probability is thus the product of the corresponding conditional probabilities. Certain of these probabilities are still poorly known so that the estimates provided here are very rough. However, producing this formula can have two important effects. First, it can help developers of gene drive technology to contemplate the risk associated to their action and consider its potential magnitude. Second, it can trigger further research for better assessment of these probabilities. A famous example of such an approach is the Drake Equation which aimed at estimating the number of active, communicative extraterrestrial civilizations in the Milky Way galaxy (Burchell 2006). The equation was written in 1961 by Frank Drake, mainly as a way to stimulate scientific dialogue. A weakness of the Drake equation is that some factors are poorly known. However, it certainly promoted numerous research studies and thoughts among scientists.

Here we examine CRISPR-based gene drives and we review the different parameters to be considered to evaluate the risk of transfer to another species. We focus on gene drives that can spread autonomously and we do not consider here binary systems, whose genetic elements are located on different chromosomes. Such binary systems have been proposed as a solution to prevent contamination of non-target populations. However, they are expected to spread less efficiently than autonomous gene drives (Esvelt *et al.* 2014; Akbari *et al.* 2015).

## Results

For a given gene drive construct to contaminate another species, six consecutive steps are required (Fig. 1). The probability of contamination can be estimated by multiplying the probability of occurrence of each event, as defined with the following Drive Risk Assessment Quantitative Estimate (DRAQUE) formula: D = (*hyb* + *transf)* × *express* × *cut* × *flank* × *immune* × *nonextinct*.

**Fig. 1.**
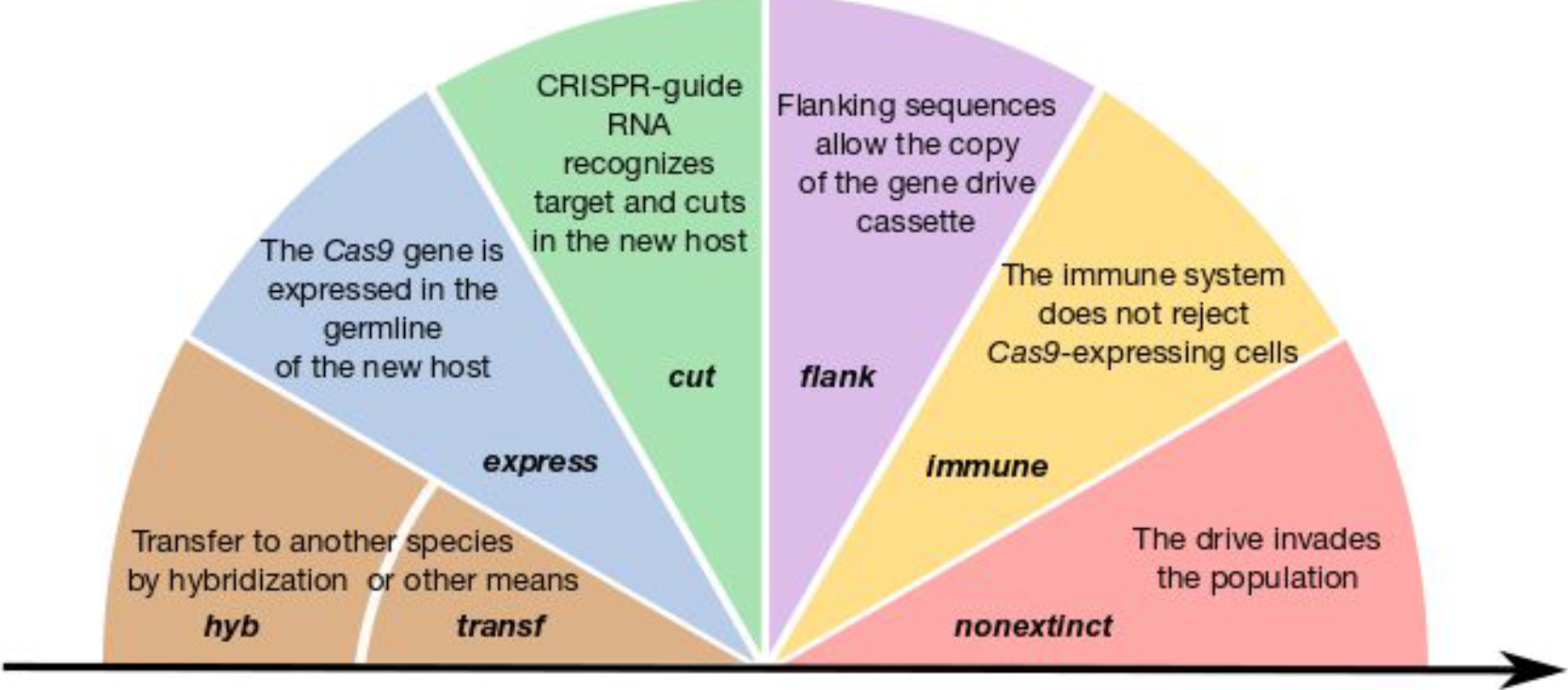
Summary of the different events whose probability must be estimated to assess the risk of gene drive contaminating another species.

The bulk probability *D* of the gene drive to contaminate a non-target species includes (i) the probability that the given piece of DNA passes from the target species to the non target species. This can occur by hybridization (*hyb*) or by other means (*transf*). Then (ii) the *Cas9* gene and the guide RNA gene have to be expressed in the new host, with probability *express*. A target sequence must then be recognized and cut by CRISPR-Cas9 (iii) with probability *cut*. The gene drive cassette should insert at the cut site (iv) with probability *flank*; the immune system (v) must not eliminate it, with probability *immune* and finally, it must not be eliminated by stochastic processes (vi) with probability *nonextinct*.

This formula could be estimated for one given non-target species. However, for ease of estimation and practical purposes, we think that it is more relevant to directly define D as the probability that at least one non-target species is contaminated..We examine below each term of the DRAQUE formula.

### Probability of hybridization between the target species and a non-target species (*hyb*)

The probability that individuals of the target species hybridize with a non-target species and produce fertile progeny has to be evaluated for each target species, as it may vary among species. We treat here two taxa for which gene drive technology is most advanced, *Drosophila* flies and *Anopheles* mosquitoes.

*Drosophila suzukii* is an invasive pest species originating from southeast Asia that invaded both America and Europe since 2008 and that attacks ripe fruits (Scott *et al.* 2018). This species is one of the most advanced systems for potential gene drive applications (Scott *et al.* 2018). Two closely related species have been described, *Drosophila pulchrella* and *Drosophila subpulchrella*; they are found in Japan, China, India and Nepal [(Takamori *et al.* 2006) and references therein]. Recent genome data suggest that hybridization probably occurred recently between *D. suzukii* and *D. subpulchrella*, which diverged about 1-9 million years ago (Conner *et al.* 2017). Furthermore, fertile hybrids between *D. suzukii* and *D. subpulchrella* have been obtained in the laboratory [(Fuyama 1983), note that *D. subpulchrella* was erroneously named *D. pulchrella* in the 1983 paper (Muto *et al.* 2018)]. Because hybridization between closely related species usually produce hybrids with reduced fitness, it can lead to reinforcement, i.e. the increase of reproductive isolation as closely related species diverge (Turelli *et al.* 2014). Considering the prevalence of reinforcement in Drosophila, one might reasonably speculate that hybridization is common in the wild. Current data thus suggests that gene drives targeting *D. suzukii* may end up in *D. subpulchrella*/*D. pulchrella* if a gene drive ever reaches areas of contact on the Asian continent.

To control malaria with gene drive, two major strategies are currently being developed. One relies on the reduction or suppression of the population of vectors and the other on genetically modifying populations of wild vectors so that they no longer transmit pathogens. The most technically advanced approach is the one conducted by the Consortium Target Malaria aiming at reducing the population of several mosquito species of the *Anopheles gambiae* complex. This complex consists of at least 8 species of the *Anopheles* genus, morphologically indistinguishable and present in sub-Saharan Africa [(Coetzee *et al.* 2013) and references therein]. Some have a large afro-tropical distribution (*An. gambiae s.s.*) while others are restricted to savannah area (*An. arabiensis*) or coastal regions (*An. merus* and *An. melas*). It also includes the species *An. quadriannulatus* that is not considered as a malaria vector. The work by Target Malaria is conducted on 3 species of this complex : *Anopheles gambiae s.s.*, *Anopheles coluzzii* and *Anopheles arabiensis* (https://targetmalaria.org/our-work/). *An. coluzzii*, formerly known as *An. gambiae* M molecular form, is defined as a separate species since 2013 (Coetzee *et al.* 2013). Among the *Anopheles gambiae* complex, the question of hybridization has been a subject of interest for geneticists and public health practitioners for decades (Fontaine *et al.* 2015). Hybridization is of high concern and interest in these mosquito populations due to the potential spread of insecticide resistance between species, and now, with the development of gene drives, of human-made transgenes.

A classical example of transfer is the geographic expansion and adaptation to arid environment by *An. gambiae* that is associated with an introgression from *An. arabiensis* into *An. gambiae*, resulting from past hybridization between the two species (Besansky *et al.* 2003; Sharakhov *et al.* 2006). The presence of the *kdr* resistance (a mutation conferring resistance against pyrethroids, insecticides largely used in impregnated bednets) in the S form of *An. gambiae* (now *An. gambiae s.s.*) and then later in the M form (now called *An. coluzzi*) has also been explained by an introgression rather than by an independant, novel mutation (Weill *et al.* 2000). This highlights the existence of gene flow between these two species. Recent studies have also highlighted high frequency of hybridization between *An. coluzzi* to *An. gambiae* in West Africa (Oliveira *et al.* 2008; Caputo *et al.* 2011; Marsden *et al.* 2011) and an asymmetric introgression from *An. coluzzi* to *An. gambiae* (Mancini *et al.* 2015). Genetic exchanges have also been detected between *An. gambiae* s.s. and *An. arabiensis* and led to the idea of particular genomic regions being more prone to cross species boundaries than others (Crawford *et al.* 2015). In the lab, introgression of a synthetic sex ratio distortion system has even been possible from *An. gambiae* to its sibling species *An. arabiensis* (Bernardini *et al.* 2019). In summary, multiple species inside the *An. gambiae* complex appear to cross-hybridize, and developers of gene drive technology may aim to develop a system able to target several of them. The higher the number of species, and thus individuals, harboring gene drive constructs, the higher the probability that the gene drive contaminates other species.

### Probability of horizontal transfer of a piece of DNA containing the gene drive cassette from the target species to a non-target species with no hybridization (*transf*)

DNA can be naturally transferred from one eukaryote species to another via so-called horizontal transfer (HT), through unknown means that may involve vectors such as viruses, microsporidia, mites or parasitoids (Houck *et al.* 1991; Gilbert *et al.* 2010, 2014; Parisot *et al.* 2014). In particular, viruses can carry over nucleic acid loads that do not directly belong to the viral specific genetic setup, but from the virus host (Gasmi *et al.* 2015; Gilbert and Cordaux 2017). Bacteria probably also constitute an important vector to transfer DNA material from one Eukarya species to another. HGT from Bacteria to Eukarya is frequent (Lacroix and Citovsky 2016) and from Bacteria to Bacteria it is the norm. This implies that environmental DNA can be rapidly contaminated by any novel construct. As cases in point, apparition of antibiotic resistance spreads rapidly from locations where it has first appeared [see e.g. (Schultz *et al.* 2017)]. HGT from Eukarya to Bacteria has been illustrated in some cases, with viruses as intermediates (Bordenstein and Bordenstein 2016). Eukaryotic genes in bacteria (EUGENs) are frequent in intracellular parasitic or symbiotic bacteria [e.g. (Hilbi *et al.* 2011)]; they may play an efficient role in gene exchange. The DNA pieces that have been observed to undergo HT are usually between 1 kb and several dozens of kb [see HTT-DB database (Dotto *et al.* 2015)], which is comparable to the size of gene drive constructs, and can go up to 150 kb in plants and animals (Inoue *et al.* 2017; Dunning *et al.* 2019).

Two types of horizontal transfer can be distinguished, horizontal transfer of transposable elements (HTT) and horizontal gene transfer (HGT). Although transposable elements (TE) also carry genes, this distinction reflects the fact that we know many cases of HTT and relatively few cases of HGT. Most HGT events that occurred in the past between distantly related species are not expected to be detected through comparison of present-day genomes because most newly inserted DNA sequences are likely to be lost by genetic drift (i.e. the probability *nonextinct* is zero). The breadth of HGT that can be approached based on comparative studies of genome sequences is thus largely underestimated. As a matter of fact, identified HGT events involve pieces of DNA that appear to increase host fitness [HGT, e.g. carotenoid synthesis genes from fungi to pea aphids (Moran and Jarvik 2010)]. Compared to other DNA sequences, TEs have particular characteristics that allow them to integrate into DNA more frequently, and they can also self-replicate in the new host after HT, so that they are more likely to be noticed.Whether the higher rate observed for HTT than for HGT is only due to the integration and replicative properties of TE is unknown. If viruses are important vectors for HT, the propensity of TE to jump from eukaryote genomes to viruses, and reversely, more frequently than random pieces of DNA (Gilbert *et al.* 2016) may also explain their higher rate of HT. Compared to TE, a gene drive cassette can also insert itself into a host genome, but its integration into DNA may be less likely and its number of copies in a genome should be lower, so that gene drives may transfer horizontally between genomes less frequently than TEs. However, if TEs are present in the vicinity of the gene drive cassette in the target species, they could end up facilitating the transfer and integration of the DNA in another host via a hitchicking process. To limit this risk, gene drives should be designed to target genomic regions that are devoid of TEs, if such regions exist in the target species.

Better than TEs, human-made gene drive constructs resemble homing endonuclease genes, which are naturally occurring mobile elements that bias their inheritance by cutting and inserting themselves at targeted sites within genomes (Burt and Koufopanou 2004; Agren and Clark 2018). A homing endonuclease gene that specifically targets the *cox1* mitochondrial gene has been transferred independently 70 times between 162 plant species within 45 different families (Sanchez-Puerta *et al.* 2008). This element is also present in several species of fungi, green algae and liverworts, highlighting extensive HT (Cho *et al.* 1998). Unfortunately, no estimate of HT rates for homing endonuclease genes is available.

Cases of HTT have been detected between extremely distantly related species (Gilbert and Feschotte 2018). For example, the *BovB* element moved at least 11 times between snakes, lizards, ruminants and marsupials (Ivancevic *et al.* 2017) and the *Mariner* element moved between nematodes, arthropods, fungi, molluscs, vertebrates and plants [(Palazzo *et al.* 2019) and references therein]. Numerous stable introductions of virus DNA into the germline of various eukaryote species have also been reported (Katzourakis and Gifford 2010; Holmes 2011; Feschotte and Gilbert 2012; Chen *et al.* 2016). In vertebrates, endogenous retroviruses (ERVs) can insert into their host genome and generate copies of themselves through germline reinfections or retrotransposition events (Gilbert and Feschotte 2018; Greenwood *et al.* 2018). A comparison of endogenous retrovirus sequences in 65 genomes identified no less than 1000 HT events between distantly related species of vertebrates (Hayward *et al.* 2015). Pan-phylogenomic analyses of endogenous retrovirus sequences revealed that rodents are a major source of retroviruses which they can transmit to other mammals such as livestock (Cui *et al.* 2015).

HT events are also ongoing now. The best known case is the worldwide invasion of *D. melanogaster* populations by a transposable element (*P*-element) originally present in the distantly-related species *D. willistoni* (Daniels *et al.* 1990; Clark and Kidwell 1997). This invasion has been carefully recorded and occurred within a few decades during the second half of the last century. Nowadays, the *P*-element is invading two other Drosophila fly species worldwide, *D. simulans*, originating from *D. melanogaster* (Hill *et al.* 2016) and *D. yakuba* (Serrato-Capuchina *et al.* 2018). *D. simulans* flies sampled before 2010 do not carry the *P*-element, indicating that this invasion is very recent. The *P*-element arose in *D. simulans* most likely through hybridization with *D. melanogaster*, but could also have occurred via the unintended escape of a few laboratory-raised *D. simulans* flies genetically engineered to carry the *D. melanogaster P*-element. In Mammals, the endogenous retrovirus sequence KoRV-A is currently observed to invade natural populations of koala (Xu and Eiden 2015) and herpes virus DNA has started to integrate into human chromosomes (Morissette and Flamand 2010).

Rough estimates of HTT rates have been obtained recently. Systematic surveys of transposable elements in complete genomes have inferred HTT events that essentially occurred during the last 10 My. They counted more than 2000 HTT events in 195 insect species (Peccoud *et al.* 2017) and more than 330 HTT in 460 arthropod species (Reiss *et al.* 2019). El Baidouri *et al.* (2014) estimated that more than 2 million HTT occurred between plant species. A study of three *Drosophila* species estimated an average rate of 0.035 HTT per transposable element family per million years between these three species, with LTR RTs and DNA transposons displaying a higher rate of 0.046 (Bartolomé *et al.* 2009). Note that the power to identify HTT events decreases as the HTT events approach the time of species divergence, so that existing quantifications are conservative.

Based on the inventory of species inhabiting Great Smoky Mountains National Park (which harbors more than 500 vertebrate species and >3900 insect species - https://dlia.org/smokies-species-tally), we can assume that a given species can be in contact with approximately 1000 distinct Eukaryote species, either directly or indirectly via viruses, bacteria or other microorganisms. As a result, the rate of HTT can be approximated to a minimum of 0.035 transfer events to at least one species per thousand of years. This estimate is conservative as it does not take into account horizontally transferred DNA that have disappeared from the recipient genome since their transfer nor HTT events that occurred right after reproductive isolation of the two lineages of interest. In addition, due to the broad geographic distribution of viruses and bacteria that can act as vectors, the probability of transfer to any species among the total estimated 10 millions in the world is probably much higher than this figure. Furthermore, there is no reason to believe that the rate of HTT is constant across time. It is possible that the number of HTT events increases under certain conditions, such as ecological stress or pervasive pathogen infections (Horváth *et al.* 2017). This stress sensitivity has been witnessed as a cause of evolution from commensalism to pathogenicity driven by an intra-organism explosion of TEs (Magaziner *et al.* 2019).

The risk of transfer to non-target species also depends on the persistence time of gene drives. For replacement/rescue drives, the drive is expected to reach all individuals of the targeted population. Then, in the long term, since there is no selective pressure to maintain a functional endonuclease, the CRISPR-cas9 cassette can eventually accumulate mutations and thus lose its self-replicating activity, and should eventually disappear from the target population. However, the persistence time of the gene drive cassette should still be relatively long, of the order of several thousands of generations, leaving time at the human scale for possible HGT. The long term presence of rescue/replacement drive constructs in the population increases the odds of transfer to non-target species. As far as we know, no strategy has been proposed to completely remove the gene drive cassette once a population has been fully targeted by a replacement drive.

For eradication/suppression drives, the target population is expected to go extinct, so that no gene drive construct is expected to remain in living organisms. Nevertheless, DNA is a very stable molecule and DNA from dead organisms can make up reservoirs of gene drive constructs. Just taking into account viruses, aquatic ecosystems typically contain virus-like particles and sediments 10^8^-10^9^ particles (Filippini and Middelboe 2007). DNA is found in all kinds of environments [(see (Hunter *et al.* 2019) for techniques allowing recovery of 1 to 500 ng of DNA per microliter of water]. Some organisms, such as *Acinetobacter baylyi* are able to take it spontaneously into their genome (Mantilla-Calderon *et al.* 2019), further amplifying and propagating DNA sequences. Subsequently, bacteria can transfer DNA into their hosts chromosomes following successful metabolic pathways, in particular via symbiotic organisms, as we have discussed previously. Gram negative bacteria are cases in point as it has been shown that they rapidly transfer antibiotic resistance to a large number of recipients (Oliveira and Reygaert 2019).

In summary, recent data indicate that DNA can transfer extensively between distantly-related species in all taxon groups. Current quantifications of the probability of transfer of a particular DNA piece to another species are rare and underestimated, and provide a probability of a minimum of 0.035 transfer events to at least one species among 1000 per thousand years. Whereas the probability of hybridization discussed above is relatively high and concerns a small number of species, the probability of horizontal transfer discussed here is relatively low but involves a larger number of potential non-target species. These non-target species can be very distant both phylogenetically and geographically, due to vectors which can be ubiquitous.

### Probability that the guide RNA and *Cas9* genes are expressed in the new host (*express*)

For the CRISPR-Cas9 gene drive system to be active in the non-targeted species, the guide RNA gene and the *Cas9* gene must be expressed in the newly-formed zygote or in the new host germline (Fig. 2). In other words, potent enhancer regions should be present in the vicinity of the two genes, and promoters should be active to drive expression in the new host (Wittkopp and Kalay 2012). Experiments swapping enhancers between species suggest that rodent sequences are active across Mammals, while fly/mosquito sequences function across Diptera and sometimes throughout insects. Human enhancers generally drive similar expression in mice [88 My divergence - all divergence times are from TimeTree (Kumar *et al.* 2017)] (Cheng *et al.* 2014). Enhancers have also been observed to drive expression in corresponding homologous organs in more distantly related species such as mice and bats [94 My divergence (Cretekos *et al.* 2008; Booker *et al.* 2016)] or even fish species and humans [465 My divergence (Yuan *et al.* 2018)]. A particular enhancer was shown to drive comparable expression across 9 species of vertebrates including fishes, lizards, snakes and mice (Kvon *et al.* 2016). In insects, several enhancers have been tested and observed to drive similar expression patterns in *Drosophila*, mosquitoes (248 My divergence) and *Tribolium* [309 My divergence (Cande *et al.* 2009; Lai *et al.* 2018)]. Several native promoters of Drosophila have been tested and they have been found to function in Lepidoptera [286 My divergence (Tamura *et al.* 2000; Imamura *et al.* 2003; Ramos *et al.* 2006)] but not in *Tribolium* (Schinko *et al.* 2010). Native mosquito enhancers are thus unlikely to function in humans (790 My divergence) or in the malaria parasite (1552 My divergence).

**Fig. 2.**
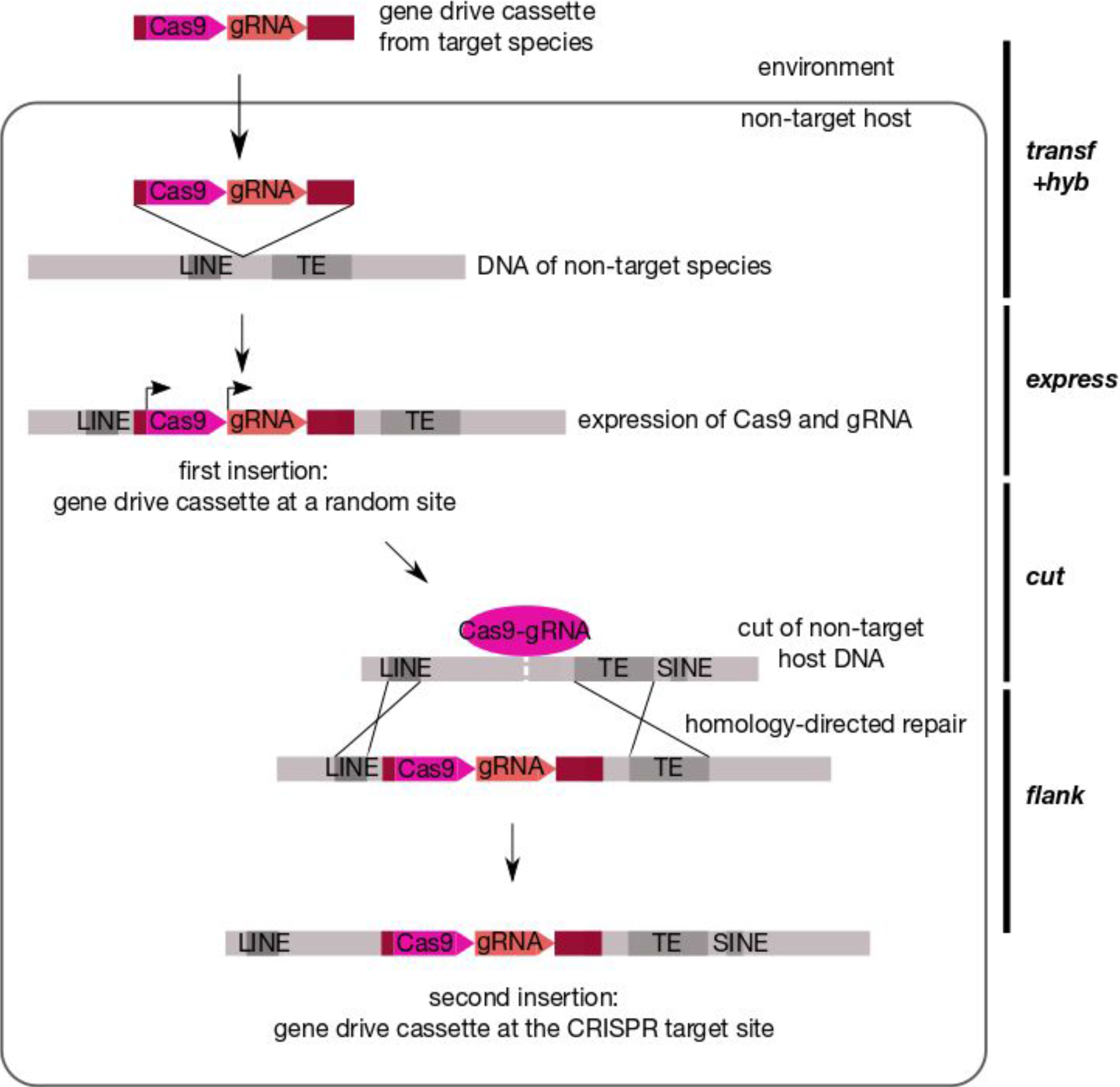
Succession of putative events leading to the insertion of a functional gene drive cassette in another distantly related non-target species (in case of horizontal transfer with no hybridization). DNA of the non-target species is indicated in gray and DNA from the target species in warm colors:pink: *Cas9* gene, orange: gRNA gene and brown: neighboring sequences. On the right are different parameters of the DRAQUE equation. In the represented scenario, one LINE (Long INterspersed Element) and one Transposable Element (TE) serve as flanking sequences for homology-directed repair. Note that other sequences present in the non-target host may also be used.

To activate expression, published gene drive constructs harbor endogenous cis-regulatory sequences of germline-specific genes for the *Cas9* gene and of constitutively expressed genes for the guide RNA genes (Table 1). Therefore, present-day constructs designed for flies or mosquitoes can be expected to drive expression in the germline across Diptera/insects, and for mice across Mammals.

**Table 1.** List of cis-regulatory sequences used in published gene drive constructs. Information is presented in the chronological order of the publications. All are endogenous cis-regulatory sequences. The genes whose cis-regulatory sequences were used to drive *Cas9* expression are expressed specifically in the germline. *U6* is a ubiquitously expressed gene encoding a small RNA involved in mRNA splicing. For *Cas9* expression in *M. musculus*, the germline-specific cis-regulatory sequences were used in combination with the Cre-loxP system and constitutive cis-regulatory sequences.

| Target Species | Cis-regulatory sequences for *Cas9* expression | Cis-regulatory sequences for guide RNA expression | Reference |
| --- | --- | --- | --- |
| *Drosophila melanogaster* | *D. melanogaster vasa* | *D. melanogaster U6:3* | (Gantz and Bier 2015) |
| *Anopheles stephensi* | *A. stephensi vasa* | *A. stephensi U6A* | (Gantz *et al.* 2015) |
| *Anopheles gambiae* | *A. gambiae vasa2* | *A. gambiae U6* | (Hammond *et al.* 2016) |
| *Drosophila melanogaster* | *D. melanogaster vasa*<br>*D. melanogaster nanos* | *D. melanogaster U6:3* | (Champer *et al.* 2017) |
| *Drosophila melanogaster* | *D. melanogaster Sry-alpha*<br>*D. melanogaster DNAPol-α180*<br>*D. melanogaster Rcd-1r* | *D. melanogaster U6:3* | (KaramiNejadRanjbar *et al.* 2018) |
| *Anopheles gambiae* | *A. gambiae zpg* | *A. gambiae U6* | (Kyrou *et al.* 2018) |
| *Mus musculus* | *M. musculus vasa*<br>*M. musculus Stra8* | *H. sapiens U6* | (Grunwald *et al.* 2019) |

Synthetic regulatory elements containing multiple adjacent binding sites for a small number of transcriptional activators have not been used so far in published gene drive constructs but they might in the future. In general, such synthetic pieces tend to be more universal than endogenous regulatory elements (Schetelig and Wimmer 2011). For example, the Drosophila-derived *3×P3* artificial element drives expression in the eyes not only in insects but also in arthropods (Pavlopoulos and Averof 2005) and planarians (Gonzalez-Estevez *et al.* 2003).

The guide RNA gene and the *Cas9* gene can also be activated by enhancers located further away, outside of the original gene drive cassette. Reciprocally, vectors used for gene therapy have been shown to misregulate endogenous genes neighboring the inserted DNA, thereby causing harmful side effects such as leukemia (Browning and Trobridge 2016). The addition of chromatin insulators into retroviral vectors, to block the activation of nearby genes, seems to be a good solution to this problem, as shown by a recent successful gene therapy (Mamcarz *et al.* 2019). To our knowledge, none of the published gene drive constructs contain insulators. To limit the activity of gene drive constructs in non-targeted species, we suggest that gene drive vectors should include insulators.

### Probability that the CRISPR-guide RNA recognizes and cuts at a DNA site in the new host (*cut*)

So far, the CRISPR-Cas9 system has been shown to cut at target sites specified by the guide RNA in all species that have been tested, including animals, plants, bacteria and parasites (Zhang 2019). We therefore consider here that the probability *cut* that the CRISPR-guide RNA recognizes and cuts at a DNA site in the new host is simply the probability that the host genome contains a site targeted by the CRISPR-guide RNA of the gene drive construct. The target site recognized by CRISPR-Cas9 is 20 nucleotides long followed by a protospacer adjacent motif (PAM), typically NGG (Zhang 2019). Importantly, studies in yeast and human cells indicate that CRISPR-Cas9 cleavage activity can still occur with three to five base pair mismatches in the 5’ end (Fu *et al.* 2013; Hsu *et al.* 2013; Roggenkamp *et al.* 2018). As a consequence, we need here to estimate the probability of finding a particular sequence of 17-19 nucleotides (plus one nucleotide that can be either base, located two bases before the 3’ end of the segment) in genomes of interest. The existence of other off-target cuts with lower sequence similarity to the on-target site is still under exploration and not entirely understood (Zhang *et al.* 2015; Gao *et al.* 2019). The number of off-target cuts tends to accumulate with longer and stronger expression of the *Cas9* gene (Kim *et al.* 2014), suggesting that the range of off-targets may differ between gene drive cassettes. We consider here only the sequences that are closely related to the on-target site and therefore provide a lower estimate of the probability *cut*.

A very rough estimate of the probability for a given sequence of 19 nucleotides to be present in a genome of a billion nucleotides can be estimated as follows. Assuming that the 4 nucleotides A, C, G and T are equiprobable, the probability of finding each sequence is 2/4^19^ (the factor 2 stands for the fact that DNA has two strands) which is about 7.10^−12^. In a genome of 3.10^9^ nucleotides such as the human haploid genome, such a sequence should be present with a probability 7.10^−12^×3.10^9^, which is approximately 2% (about 30% for a sequence of 17 nucleotides). Note that this is a very crude estimate. It could be higher if one took into account the fact that the DNA contents of AT and GC are different. Also, it does not take into account that an extant DNA sequence is never random. The non random character of genomes is illustrated by the omnipresence of repeated sequences (for example in the human genome the repeats Alu, SINE, LINE etc.) so that if a given sequence is present in a genome, it is likely to be present more than once. Overall it appeared that genomes behave as n-plications of a core set of sequences followed by reduction. This has been observed in yeasts (Escalera-Fanjul *et al.* 2018), plants (Clark and Donoghue 2018a) and animals (Hermansen *et al.* 2016).

The fact that the target sequence is not an arbitrary sequence can increase or decrease the probability according to whether the bias for the sequence of interest and the bias for the sequences in the genome both go in the same direction or not. The probability *cut* would be lower if the existence of repeated sequences was taken into account and if the targeted sequence did not match some of the repeated sequences. By contrast, it would be higher if the sequence matched the “style” of the DNA of a particular organism, i.e. its non random content in particular motifs (Fertil *et al.* 2005). Different mechanisms can lead to the occurrence of repeated sequences in the genome (gene conversion, unequal gene exchanges, transposition, etc.); they have been grouped under the name of “molecular drive” (Dover 1982). “Molecular drive” is likely to be widespread, as a sizeable proportion of individuals carry local duplications of any sequence of the genome. Furthermore, some conserved repeats may maintain the coexistence of stable rearrangements in some species (Smalec *et al.* 2019), increasing the possibility of unexpected cuts in certain species. All these features may considerably impact the probability of cuts in particular genomes. If the targeted site is present in multiple copies, there is a risk for the gene drive construct to spread across the entire genome. We urge the researchers developing gene drive constructs to make sure that they choose a target site that is very distinct from sequences such as retroelements, LINE, SINE elements that are present in large quantities in genomes [see (Breitwieser *et al.* 2019) for an updated view of repeats in the human genome].

A recently proposed gene drive that aims at sterilizing mosquito females has the following target site, GTTTAACACAGGTCAAGCGGNGG, which is a highly conserved sequence within the *Doublesex* gene, displaying a 3’ terminal end that contains either a repeated CGG triplet or variant of it. We should note that CGG repeats are quite frequent in genomes (Pan *et al.* 2018; Rabeh *et al.* 2018). The 23-bp sequence is present in 7 species of *Anopheles* mosquitoes, and with one mismatch in at least 6 additional *Anopheles* species (Kyrou *et al.* 2018). Sequencing of 765 wild-caught *Anopheles gambiae* mosquitoes identified only one single nucleotide variant within this sequence, and this variant was still permissive to gene drive. Our BLAST searches for this target sequence and for the one used in the published mouse gene drive found fragments of up to 20 bases identical to the 3’ part of the sequence in several genomes, belonging to all three domains of life (Table S1). While a target sequence which accommodates little nucleotide variation such as the *Doublesex* locus can be useful to prevent gene drive resistance, it is then also associated with an increased risk of spread to non-target species. This trade-off exists not only for the *Anopheles gambiae* mosquito complex but for any species targeted by a gene drive system.

### Probability that the gene drive cassette inserts near the targeted site (*flank*)

The gene drive cassette is designed to bias its transmission by copying itself on the paired chromosome, so that heterozygous individuals carrying initially one copy of the gene drive cassette end up with two copies in their germline cells (Esvelt *et al.* 2014). This homing process occurs through homology-directed repair, using homology arms flanking both the gene drive cassette and the guide RNA target site. Therefore, for the gene drive to be active in non-target species, the DNA containing the *Cas9* gene and the *gRNA* gene should be inserted at the guide RNA target site. *Flank* is the probability that the gene drive cassette lands up at the guide RNA target site.

In case of hybridization, the non-target species being closely related to the species carrying the initial gene drive, the genomic regions harboring the gene drive cassette are expected to be comparable, so that the probability *flank* should be close to 1.

In case of horizontal transfer to a distantly related species with no hybridization, the gene drive cassette is likely to insert at a random site in the genome. In this case, *flank* is the probability that it moves from this initial position to a site targeted by its guide RNA (Fig. 2). There are several ways such a transposition can happen. Double-strand DNA breaks, such as the one created at the guide RNA target site by the CRISPR system, are known to induce recombination and various DNA repair mechanisms (Jasin 1996; Hartlerode and Scully 2009).

First, the gene drive cassette may move to the guide RNA target site via homology-directed repair (Fig. 2). Such phenomenon has been observed with gene drive constructs that were not inserted initially at the target site but at another position in the genome (Gantz and Bier 2015; Guichard *et al.* 2019). The probability of such an event will depend on the length of the flanking homology arms, their percentage of identity,the length of the DNA sequence in between and the position of the donor sequence relative to the cut site (Wang *et al.* 2017). Unfortunately, as far as we know, no extensive survey of these parameters has been done with respect to homology-dependent repair using another chromosomal locus as template. A gene drive cassette of 21 kb was found to spread effectively in mosquitoes (Gantz *et al.* 2015), suggesting that relatively large pieces of DNA can be inserted via homology-directed repair. In any case, larger inserts tend to show lower efficiency of recombination (Li *et al.* 2014). Gene drive constructs published to date have used flanking sequences of about 1kb (Gantz and Bier 2015; Gantz *et al.* 2015; Grunwald *et al.* 2019). When linear double stranded DNA is used as a template for repair, efficient targeted genome integration can be obtained using flanking sequences that are only 50-bp long in mammalian cells (Li *et al.* 2014; Wierson *et al.* 2018) and even 20-40-bp long in zebrafish (Auer and Del Bene 2014; Hisano *et al.* 2015; Wierson *et al.* 2018), *Xenopus laevis* (Nakade *et al.* 2014), *Bombyx mori* (Nakade *et al.* 2014) or the nematode *C. elegans* (Paix *et al.* 2014). Whether such short sequences could also favor transposition of the gene drive cassette to the guide RNA target site remains unknown. If repeat sequences exist both near the initial insertion site of the gene drive and the guide RNA site, they may facilitate such transposition (Fig. 2). Evaluating the parameter *flank* thus requires an assessment of the distribution of repeats in potential non-target species.

As emphasized by Salzberg and coworkers, the human genome sequence, while far more complete than most animal genomes, is still made of 473 scaffolds and comprizes 875 gaps (Breitwieser *et al.* 2019). As expected, the gaps encompass regions with a variety of repeats, some of them still poorly characterized in particular in centromeric and pericentromeric regions. This precludes accurate analysis of their distribution, and the situation is even worse for other genomes. For instance, the transposon-derived Alu repeats (~300 bp long), that are present in primate genomes at more than one million copies, are widely variable, within and among chromosomes (Grover *et al.* 2004). Interestingly this distribution is biased towards proximity of protein-coding gene regulatory regions (Lavi and Carmel 2018). In addition to our somewhat limited knowledge of the distribution of repeats in mammalian genomes we have to consider that when a species is represented by a large number of individuals there are many copy number variants, in particular in repeated regions (Monlong *et al.* 2018). This is an unavoidable consequence of involvement of satellite DNA in recombination at heterologous sites, a process which is still a matter of speculation (Satović *et al.* 2016).

Second, the gene drive cassette may move to the guide RNA target site via a transposable element. It is well established that transposable elements play a considerable role in displacement of genes or regulatory regions across genomes (Chen and Yang 2017). As a matter of fact TEs are recognized as a frequent cause of genetic diseases [see for example (Larsen *et al.* 2018; Song *et al.* 2018) and references therein].

The events discussed above may seem extremely rare, but they do not have to occur right after the gene drive inserted in a genome. The gene drive cassette may stay dormant for a few generations, and there can be many trials and errors in various individuals before an insertion occurs at the guide RNA target site. Of note, gene drive constructs containing several guide RNAs (Esvelt *et al.* 2014) increase the chance that the drive moves to a targeted site.

### Probability that the immune system does not eliminate Cas9-expressing cells (*immune*)

The Cas9 protein is derived from the bacteria *Streptococcus pyogenes* (Zhang 2019) and can trigger an immune response in mice (Chew *et al.* 2016; Chew 2018). During the gene drive multiplication process, germline cells produce Cas9 proteins to cut the DNA and insert the gene drive cassette (Fig. 2). These germline cells may thus present Cas9 fragments on their surface.

In insects, the immune system is not known to recognize novel foreign molecules (Lemaitre and Hoffmann 2007), hence the presence of Cas9 is unlikely to trigger an immune response. However, double stranded RNAs larger than 30bp can be recognized by Dicer2 and activate the RNA interference pathway, leading to their degradation (Elbashir *et al.* 2001; Gammon and Mello 2015). Guide RNAs are about 100 nucleotide long including the target site and they contain hairpins shorter than 15bp (Bassett and Liu 2014), so they should not be recognized by Dicer2. In summary, present knowledge thus suggest that gene drives would not be hampered by the immune system in insects.

In vertebrates, if Cas9 proteins accumulate in somatic tissues at a late stage during development due to leakage of the cis-regulatory regions controlling *Cas9* gene expression, Cas9 fragments may be recognized as foreign molecules and trigger an adaptive immune T cell response, leading to the potential elimination of the Cas9-expressing cells and a probable decrease in fitness of the individual carrying the gene drive (Chew 2018). The expression of *Cas9* at an early stage of development may also activate an immune response if the gene drive carrier has herited anti-Cas9 antibodies from its mother, as maternal immunoglobulins G have been shown to cross the placental barrier and the intestine, and to be maintained for a long time in the fetus after birth (Madani and Heiner 1989; Roopenian and Akilesh 2007). However, testes - and maybe ovaries - appear to readily accept foreign antigens without the induction of an immune response in several mammals (Simpson 2006; Mellor and Munn 2008; Li *et al.* 2012), so that germline cells expressing *Cas9* may not be eliminated by the immune system. Furthermore, guide RNAs produced from gene drive constructs are not expected to elicit an immune response in vertebrates, as they do not carry 5’-triphosphate ends (Kim *et al.* 2018; Wienert *et al.* 2018). Clearly, our knowledge in immunology is presently too sparse to anticipate how gene drive systems will interact with the immune system in vertebrates.

*S. pyogenes* is a facultative pathogenic bacteria mostly restricted to humans, with about 10-20% of the population being asymptomatic carriers (Shaikh *et al.* 2010; Roberts *et al.* 2012). It is thus no surprise that recent investigations have found anti-Cas9 antibodies and Cas9-reactive T cells in several human populations (Charlesworth *et al.* 2019; Wagner *et al.* 2019). It is thus possible that certain humans are immune to gene drives, and future studies will undoubtedly shed light on this question. The presence of *S. pyogenes* has also been documented in a few animals such as macaques, mice, dogs, hedgehogs, rabbits and sheep [(Vela *et al.* 2017; Chen *et al.* 2019) and references therein]. Whether these vertebrates may also be immune to gene drives remains to be investigated.

So far, all published gene drive constructs have used Cas9 from *S. pyogenes* (SpCas9). Other Cas proteins with activity similar to SpCas9 are available (Zhang 2019) and could potentially be used for gene drive technology. As they are derived from other types of bacteria, their immunogenicity and their associated probability *immune* would have to be specifically assessed.

### Probability of non-extinction of the drive (*nonextinct*)

Once the drive successfully introduced into the genome, its fate will depend on its ability to distort its segregation, on chance and on the probability that the non-target population evolve resistance to the gene drive. For simplicity, we treat here the probability that a drive, initially present in a single individual or so, is not eliminated immediately by mere chance due to sampling effect and reach significant numbers of individuals in the non-target population. Even if an allele manages to be present in more than one half of the gametes from heterozygous individuals, it can still disappear rapidly due to random processes. If not, then it can invade the population. This process has been modelled as a branching process by Bienaymé in 1845 (Bienaymé 1845), actually published by Cournot (1847), and Watson and Galton (1875), see (Kendall 1975; Bru 1991). The probability of extinction, starting with one replicator, is the lowest root of equation *G*(*x*) = *x*, where *G*(*x*) is the generating function of the law describing the number of copies left per generation by one replicator. Here, it is relevant to assume the distribution to follow a Poisson law. If the population as a whole is stable, each “normal” gene leaves on average one copy of itself in the next generation, *i.e.* on average 1/2 through male gametes and 1/2 through female ones. The drive will then leave a number of copies of itself equal to the sum of the proportions which it represents in male (λ_m_) and in female (λ_f_) gametes : λ = λ*m* + λ*f*. Typically, λ*m* and λ*f* are around 0.7 – 0.9 for gene drives tested in mice and insects (Gantz and Bier 2015; Gantz *et al.* 2015; Hammond *et al.* 2016; KaramiNejadRanjbar *et al.* 2018; Kyrou *et al.* 2018; Grunwald *et al.* 2019) so that a reasonable estimate of λ is 1.4 – 1.8. According to the “Bienaymé, Galton, Watson” model, the extinction probability for a Poisson law of parameter λ is such that *G*(*x*) = *e* ^λ(*x*−1)^ = *x*. With λ = 1.4 (resp. 1.8), it gives *x* ≃ 0.5 (resp. 0.27) and the probability of invasion (1 − *x*) is thus around 0.5-0.7. For the dynamics of gene drive alleles once they are introduced in a substantial amount in a targeted population, several stochastic and deterministic models have been composed, which take into account possible costs of harboring a gene drive as well as appearance of resistance to gene drive (Marshall 2009; Unckless *et al.* 2015).

## Discussion

Based on our examination of the seven parameters of the DRAQUE Equation (Table 2), the probability that a gene drive transfers to another species can have values ranging from 0 to 0.5 per year in the worst-case scenario (one transfer event occurring per year, the guide RNA site is present in the non-target genome, the non-target species has high levels of homologous recombination). Our current estimate of the overall risk (Table 2) remains nevertheless very crude, and asks for further studies to refine this estimate.

**Table 2.**
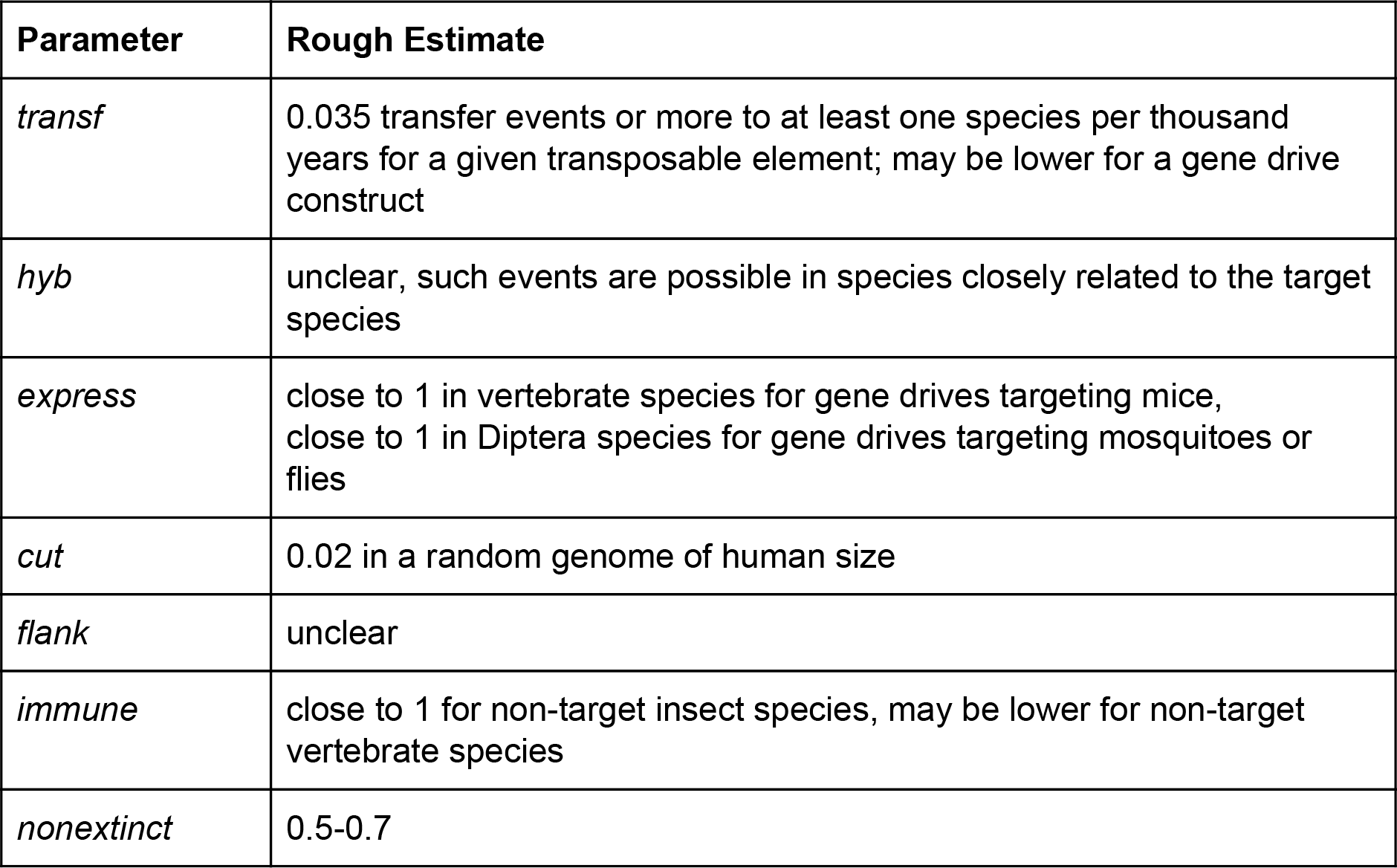
Overview of the various parameters of the DRAQUE Equation.

| Parameter | Rough Estimate |
| --- | --- |
| *transf* | 0.035 transfer events or more to at least one species per thousand years for a given transposable element; may be lower for a gene drive construct |
| *hyb* | unclear, such events are possible in species closely related to the target species |
| *express* | close to 1 in vertebrate species for gene drives targeting mice, close to 1 in Diptera species for gene drives targeting mosquitoes or flies |
| *cut* | 0.02 in a random genome of human size |
| *flank* | unclear |
| *immune* | close to 1 for non-target insect species, may be lower for non-target vertebrate species |
| *nonextinct* | 0.5-0.7 |

We did not treat here the risk of contamination of another population, within the targeted species, but the same formula could be used in principle for this case. Furthermore, our equation does not take into account the phenotypic effect of the drive on the contaminated species. A drive may display no phenotypic effects in the non-target species that it invaded. If the gene drive was designed to eliminate a target population, then it is more likely to eliminate the non-target species. The drive may also have unexpected effects, for example mosaicism due to perdurance of Cas9 expression (Guichard *et al.* 2019).

To prevent contamination, several containment strategies for laboratory experiments have been proposed (Benedict *et al.* 2008; Akbari *et al.* 2015; National Academies of Sciences and Medicine 2016) as well as gene drives split in two different constructs (Benedict *et al.* 2008; DiCarlo *et al.* 2015) and synthetic target sites (Champer *et al.* 2019). Based on our survey of the various parameters, we suggest further design strategies to minimize the risk of transfer to a non-target species: the addition of insulators (see above), and the choice of a guide RNA target site that is not close to the centromere, to avoid rearrangements and increased probability of creating an active gene drive in a non-target species.

Here we treated probabilities for a standard gene drive construct, but the risk should be estimated for each particular gene drive construct. New types of self-limiting gene drives have been proposed in recent years to try to limit the spread of gene drives spatially or temporally: toxin-antidote systems including Medea (Buchman *et al.* 2018), CleaveR (Oberhofer *et al.* 2019), Killer-Rescue (K-R) (Gould *et al.* 2008; Webster *et al.* 2019), one or two-locus underdominance (Dhole *et al.* 2018) and Daisy-Chain drives (Noble *et al.* 2019). The DRAQUE parameters would have to be evaluated specifically for these particular cases.

We have restricted our evaluation of the probability of accidental spread of gene drive constructs to genomes considered as fairly stable entities. However, genomes are dynamic structures. The process of gene duplication (or even n-plication) is commonplace (Clark and Donoghue 2018b; Harari *et al.* 2018; Moriyama and Koshiba-Takeuchi 2018). Local amplification of sequences is also very frequent (Liu *et al.* 2019; Traynor *et al.* 2019) and this may increase the probability of accidents to a further unknown level.

The DRAQUE Formula does not cover all the risks associated with gene drive. In addition to the risk of transfer to another species, gene drive designed to eliminate a target population may have additional ecological consequences that are not treated here.

Living with highly evolved technologies entails high risks for individuals and societies. Here we have attempted to evaluate circumstances where risks could be identified, but this assumes that we are aware of all the natural processes coupled with the technologies of interest. Besides the technology itself —which can be properly monitored and steered— there is an additional risk that is seldom taken into account, the risk derived from the way the organizations that implement the technologies manage them (Perrow 2011). We have not tackled this question here, but it is an obvious place of much concern. The way novel biological technologies have been used recently —see modification of human babies with deletion of a surface cell receptor (Sand *et al.* 2019)— should remind us that rogue or inconscient scientists may use gene drive approaches without proper risk assessment.

Overall, our study reveals that there is a need for more detailed investigations of the different factors influencing the probability of a gene drive contaminating another species before any release in the wild population is ever considered. We hope that our paper will trigger discussions and progress in the ethics of gene drive implementation.

## Acknowlegements

We deeply thank Florence Débarre, Clément Gilbert, Arnaud Martin and Nicolas Rode for their comments on a previous draft, and Christophe Antoniewski, Sylvain Charlat, Jean David, Scott Gilbert, Frank Jiggins, Bruno Lemaitre, Olivier Panaud, Thomas Pradeu, Benjamin Prud’homme, Michael Turelli and Mylène Weill for interesting discussions and useful information.

## Abbreviations

CRISPR: Clustered Regularly Interspaced Short Palindromic Repeats
DRAQUE: Drive Risk Assessment Quantitative Estimate
HGT: horizontal gene transfer
HTT: horizontal transfer of transposable element
TE: transposable element

---

**Table S1.**
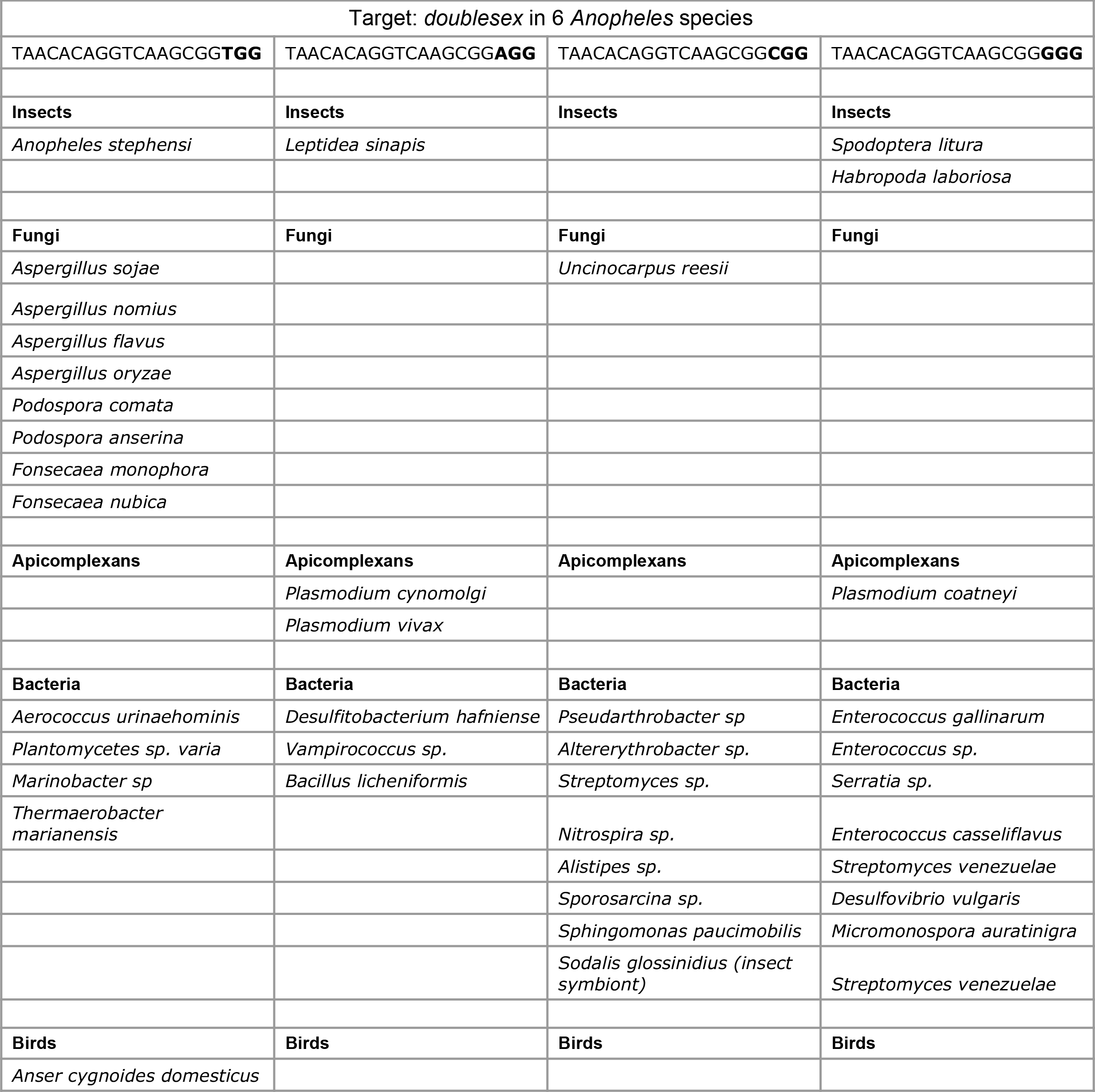
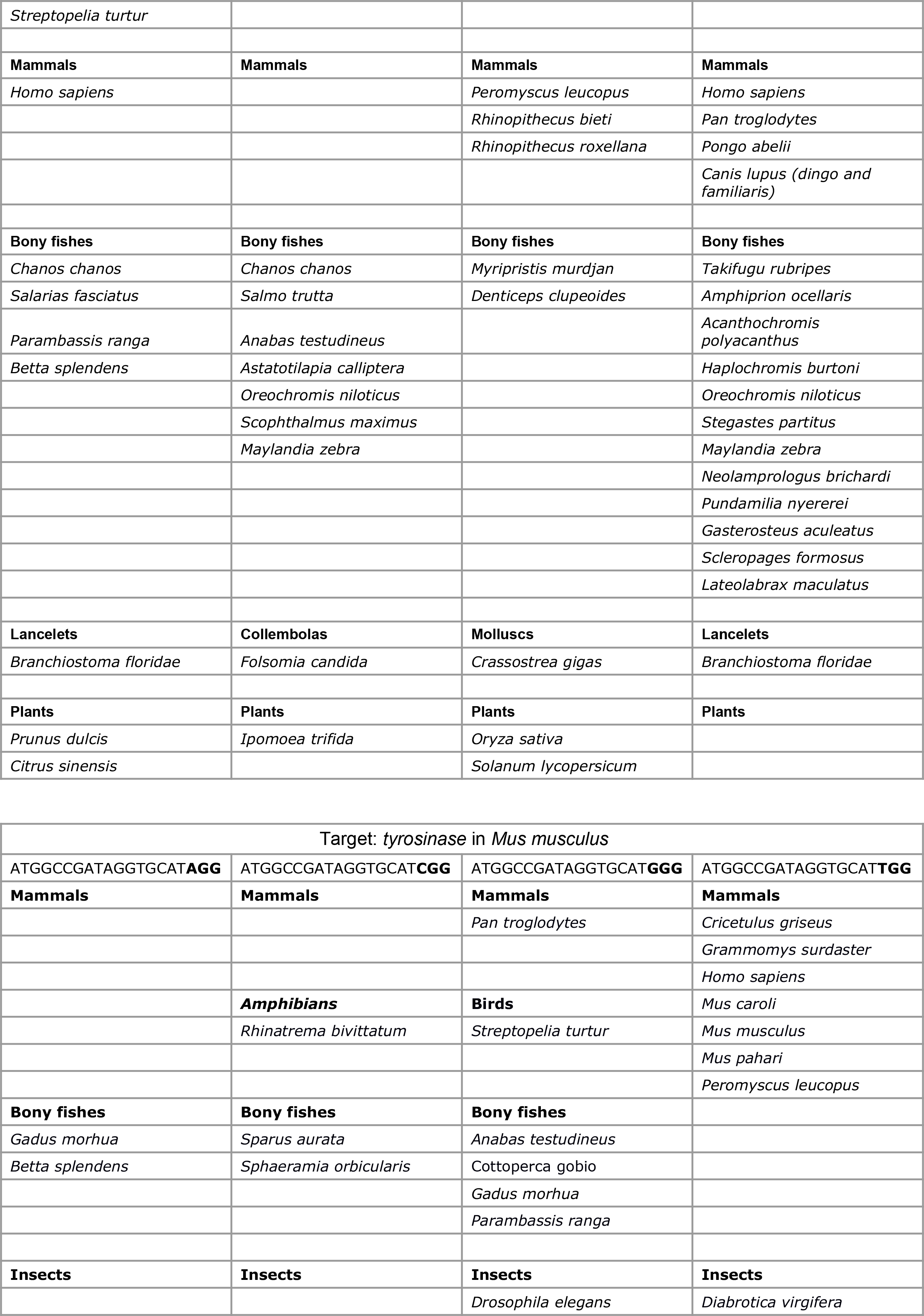
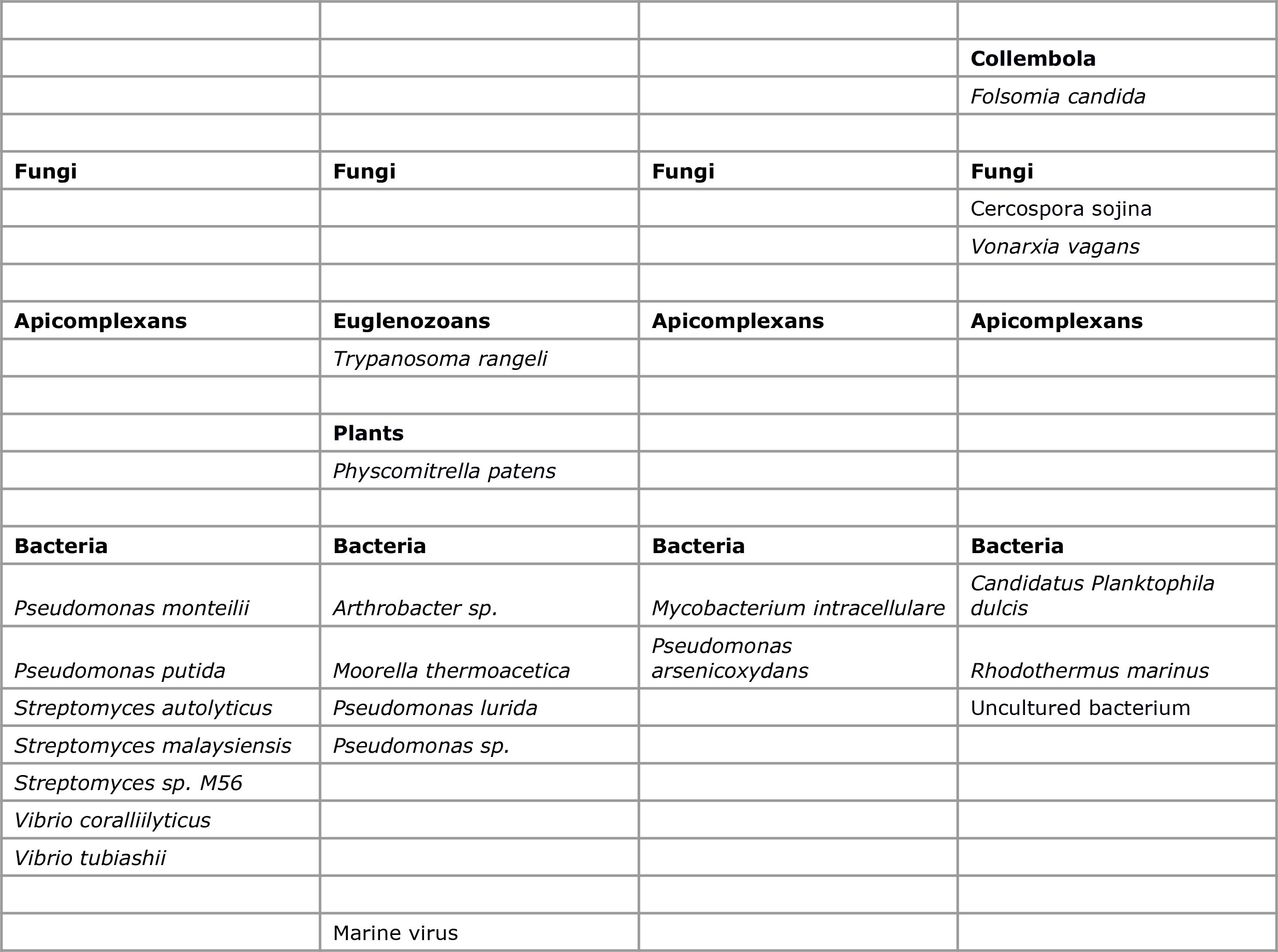
Results of BLAST searches for target sequences of two gene drive constructs published previously (Kyrou *et al.* 2018; Grunwald *et al.* 2019). The BLAST search was performed using 20-nucleotide-long queries on the nr database with default parameters on https://blast.ncbi.nlm.nih.gov/Blast.cgi?PROGRAM=blastn&PAGE_TYPE=BlastSearch&LINK_LOC=blasthome. Note that the BLAST algorithm does not retrieve results when the query sequence contains an N at the antepenultimate position. In consequence, the “N” in “NGG” was replaced by each of the four nucleotide letters and four BLAST searches were performed for each target sequence. We examined only the first 100 hit sequences and selected here the ones with at least 16/20 nucleotide sites identical to the query and harbouring the protospacer adjacent motif (PAM) NGG. Hit sequences are shown by taxonomic group for each query sequence. Note that sequences are not always deposited in the nr database (for example the *Anopheles* sequences were not retrieved by our BLAST searches), so that this table does not reflect the full extent of all published sequence data. Note also that the target sequence for the tyrosinase locus is incorrectly labelled in Extended Data Figure 2 of Grunwald *et al.* 2019 (H. Grunwald, personal communication).

